# Perinatal outcomes, maternal age, parity and fetal sex – searching for the “optimal” maternal age

**DOI:** 10.1101/365502

**Authors:** Šárka Kaňková, Jaroslav Flegr, Radim Kuba, Pavel Calda

## Abstract

**Background:** Maternal age, parity and fetal sex are each known to affect obstetric and birth outcomes. The objective of the present study was to investigate the influence of the combination of maternal age, parity and fetal sex on outcomes of pregnancies._The aim of the study was to analyze the influence of maternal age on perinatal outcomes in both primiparous and multiparous women with different fetal sex.

**Methods:** The cross-sectional study was performed on data from 11,780 women, who have given birth at the General University Hospital in Prague, Czech Republic between 2008-2012.

**Results:** Maternal age significantly (*P*<0.01) influenced pregnancy weight gain, duration of pregnancy, birth weight and birth length. Primiparous women with female newborns aged ≤19 had higher rates of preterm delivery than comparable women 20-39 (*P*=0.012). Similarly, women with male newborns aged ≥40 had higher rates of preterm delivery than comparable women 20-39 (*P*=0.003). Women aged ≤24 expressed higher rates of low birth weight than women aged >24 (*P*<0.001), regardless of parity and fetal sex. The older (≥35) primiparous women with male newborns expressed a higher incidence of macrosomia (*P*=0.021) compared to other groups of women. The probability of caesarean delivery increased with age (*P*<0.001) and it was significantly affected by the parity and sex of the newborn with higher rates of caesarean section in primiparous women as well as in mothers carrying male fetuses.

**Conclusions:** Our results indicate that “optimal” maternal age without obstetrics and birth complications is 25-34 years, older age is associated with increased complications with a male fetus, especially in primiparous women. Our data suggests that not just the age of women, but the combination of age, parity, and fetal sex should be taken into consideration during assessment of health risks of pregnancy.

## Introduction

During the last decades, there has been a clear trend in higher-income countries towards delaying childbirth to later reproductive years. In contrast to this broader population trend, rates of teenage motherhood have remained relatively stable at around 10–40 births per 1,000 per year [1–3].

Several publications have reviewed perinatal outcome either among women of advanced maternal age [4–10] or among young maternal age [5, 11–14]. Age 35 years or older in pregnant women was associated with an increased risk of intrauterine fetal death, pregnancy-induced hypertension, gestational diabetes, delivery by caesarean [8, 9, 15] and maternal near miss, maternal death and several maternal outcomes [6]. Preterm birth, gestational diabetes, and preeclampsia were more common among women 40 years or older [10]. Also, it is well acknowledged that teenage pregnancies are at increased risk for adverse birth outcomes like stillbirth, preterm birth, neonatal death, and low birth weight. [5, 11–14].

Furthermore, most published studies categorized maternal age to analyze the association. Weng et al. [5] explored nationwide population-based data of over 2 million births by each year of maternal age to comprehensively analyze the adverse birth outcomes. The data suggest that the optimal maternal ages to minimize adverse birth outcomes are 26–30 years [5]. Also, fetal sex is known to affect outcomes of pregnancies. Women carrying male fetuses were reported to be at increased risk of preterm delivery, preeclampsia, fetal distress, labor dystocia, operative delivery, and perinatal mortality [16–21]. Hyperemesis gravidarum and hypertension-related growth retardation were significantly more common in pregnancies with female fetuses [22–25]. The influence of the combination of maternal age and fetal sex on pregnancy outcomes in term and post-term singleton pregnancies were also studied by Weissmann-Brenner et al. [26].

As far as we know, the combined impact of maternal age, parity and fetal sex on the outcomes of pregnancy has not been evaluated yet. Therefore, the aim of our study was to analyze the influence of maternal age (six age categories) on perinatal outcomes in both primiparous and multiparous women with different fetal sex. Based on the combination of maternal age, parity and fetal sex we would like to find the risk groups of mothers for adverse fetal outcomes.

## Material and Methods

### Patients

The study was designed as a cross-sectional study. The main data set covered women, who have given birth in the General University Hospital in Prague, Czech Republic between 2008–2012. Clinical records comprised maternal age, maternal body weight before pregnancy, number of previous deliveries, maternal weight before conception, pregnancy weight gain, duration of pregnancy, mode of delivery, birth weight, birth length and sex of the newborn. We studied the associations of these independent variables with following adverse binary outcomes: preterm delivery (delivery before 37 weeks of gestation; yes/no), low birth weight (birth weight < 2500g; yes/no), macrosomia (birth weight > 4500g; yes/no), and caesarean delivery (yes/no). The women those giving birth to twins were excluded from the analyses. During the whole study, we worked with an anonymized data set. The project was approved by IRB Faculty of Science, Charles University (No. 2017/26) (Etická komise pro práci s lidmi a lidským materiálem Příodovědecké fakulty Univerzity Karlovy).

### Statistics

The statistical program Statistica 10.0 was used for all statistical testing. Participants, both primiparous and multiparous women, were divided into six age categories ≤ 19 years, 20–24 years, 25–29 years, 30–34 years 35–39 years and ≥ 40 years by maternal age at conception. The influence of maternal age on pregnancy weight gain, duration of pregnancy, birth weight and birth length was analyzed using ANCOVA, General Linear Model (GLM). Because of non-monotone effects of age on the output variables, the variable age was analyzed as a categorical predictor variable. The variables maternal weight before conception, pregnancy weight gain and duration of pregnancy were successively added to the statistical model as continuous predictor variables. Association between age and output binary variables preterm delivery, low birth weight, macrosomia, and caesarean section was analyzed with the logistic regression, separately for four groups of women – primiparous/multiparous women with male/female newborns. For some women, certain data were not available and therefore numbers of women varied between analyses.

## Results

The total data set contained records of 11,780 women. Table 1 shows age group-, sex of newborn- and parity-stratification of the data set. The influence of maternal age (six categories ≤ 19 years, 20–24 years, 25–29 years, 30–34 years 35–39 years and ≥ 40 years) and weight before conception, pregnancy weight gain, duration of pregnancy on pregnancy weight gain, duration of pregnancy, birth weight and birth length was evaluated using GLM, see Fig. 1. Table 2 shows results of the statistical analyses (*P*-values and effect sizes for analyzing variables) for all models.

**Fig 1.**
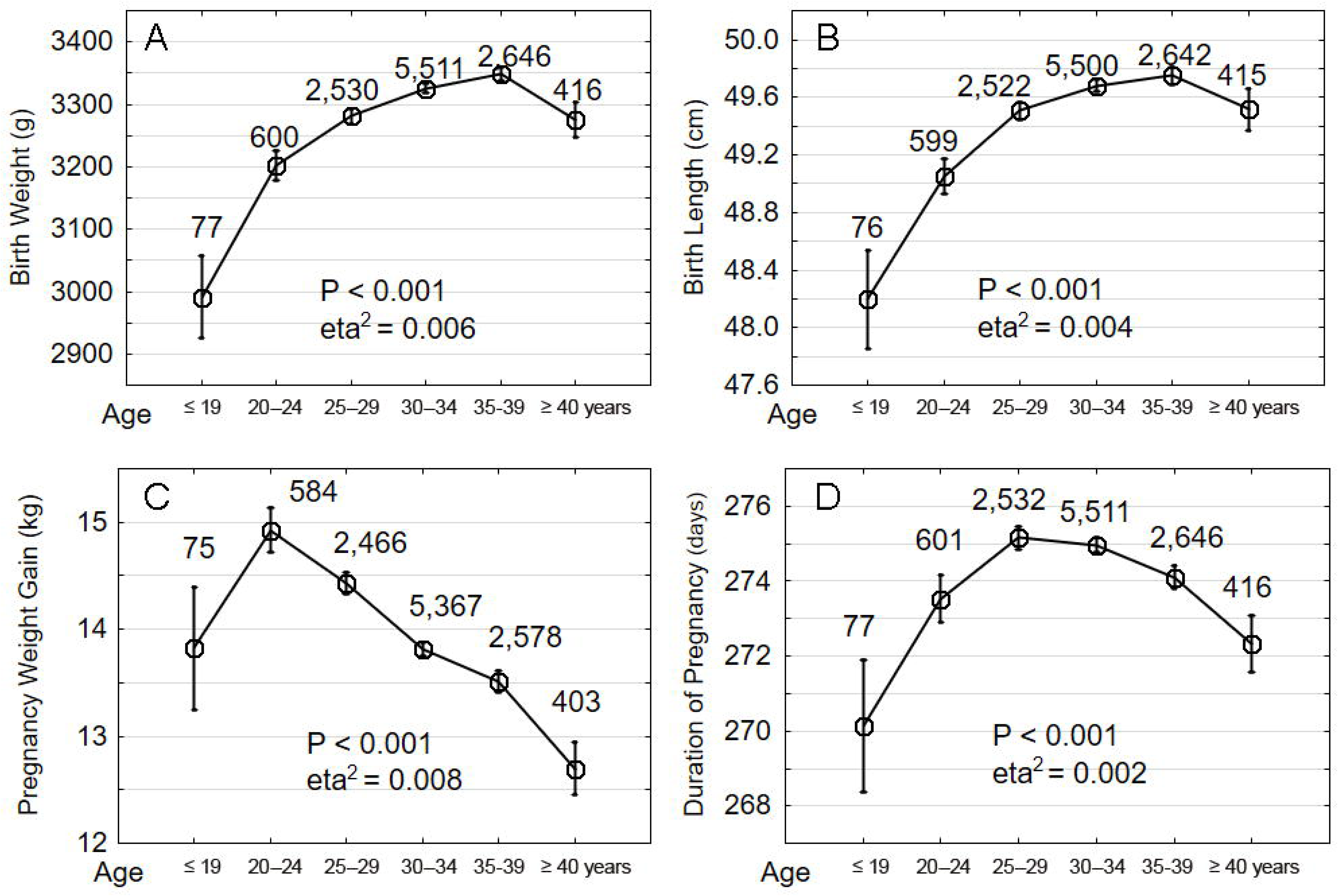
Comparison of the birth weight (A), birth length (B), pregnancy weight gain (C) and duration of pregnancy (D) in women with six age groups. The x-axis shows the age groups by maternal age at conception: 1: ≤19 years, 2: 20–24 years, 3: 25–29 years, 4: 30–34 years, 5: 35–39 years and 6: ≥ 40 years. Vertical bars denote +/- standard errors. The numbers above the boxes show the number of women in a particular category.

**Table 1:** Composition of the data set.

| Age groups | Primiparous women |  | Multiparous women |  |
| --- | --- | --- | --- | --- |
| years | male | female | male | female |
| Total | 3,660 | 3,457 | 2,368 | 2,295 |
| ≤19 | 22 | 22 | 16 | 17 |
| 20-24 | 196 | 163 | 133 | 108 |
| 25-29 | 811 | 781 | 465 | 472 |
| 30-34 | 1721 | 1632 | 1105 | 1054 |
| 35-39 | 784 | 740 | 569 | 553 |
| ≥40 | 126 | 119 | 80 | 91 |

Maternal age, parity, fetal sex and adverse fetal outcomes
age group-, sex of newborn- and parity- stratification

**Table 2.**
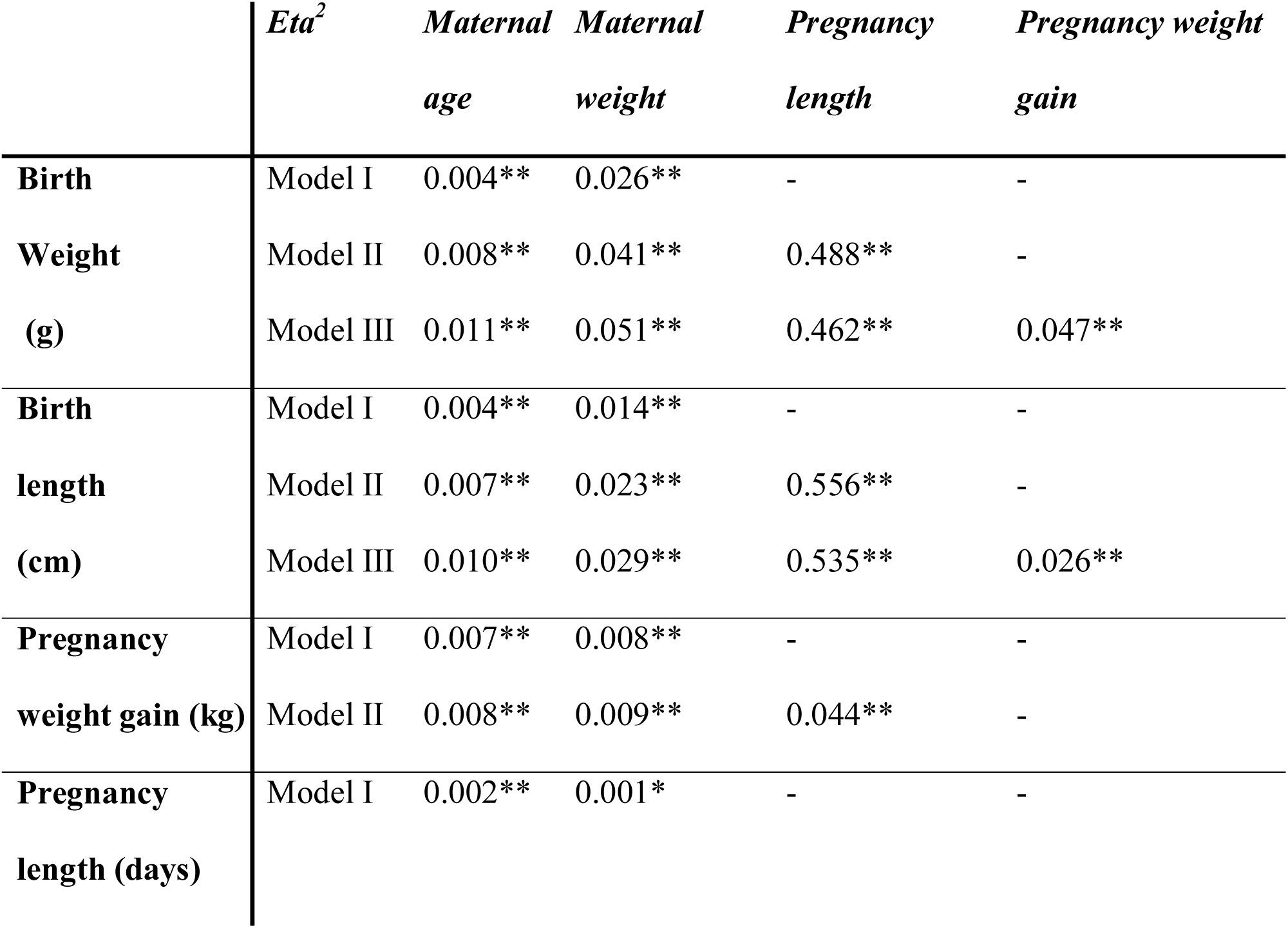
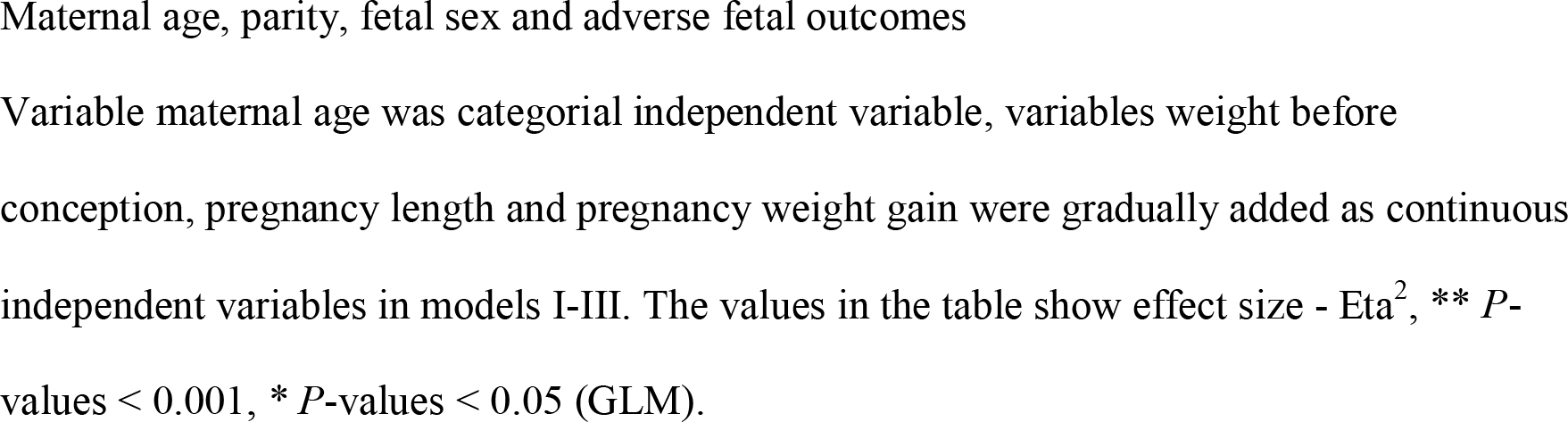
The influence of maternal age (six age categories: ≤ 19 years, 20–24 years, 25–29 years, 30–34 years 35–39 years and ≥ 40 years) on birth weight, pregnancy length, pregnancy weight gain and birth length in three models (GLM).

|  | *Eta<sup>2</sup>* | *Maternal age* | *Maternal weight* | *Pregnancy length* | *Pregnancy weight gain* |
| --- | --- | --- | --- | --- | --- |
| **Birth** | Model I | 0.004** | 0.026** | - | - |
| **Weight** | Model II | 0.008** | 0.041** | 0.488** | - |
| **(g)** | Model III | 0.011** | 0.051** | 0.462** | 0.047** |
| **Birth** | Model I | 0.004** | 0.014** | - | - |
| **length** | Model II | 0.007** | 0.023** | 0.556** | - |
| **(cm)** | Model III | 0.010** | 0.029** | 0.535** | 0.026** |
| **Pregnancy** | Model I | 0.007** | 0.008** | - | - |
| **weight gain (kg)** | Model II | 0.008** | 0.009** | 0.044** | - |
| **Pregnancy** | Model I | 0.002** | 0.001* | - | - |
| **length (days)** |  |  |  |  |  |

Maternal age, parity, fetal sex and adverse fetal outcomes
Variable maternal age was categorial independent variable, variables weight before conception, pregnancy length and pregnancy weight gain were gradually added as continuous independent variables in models I-III. The values in the table show effect size - $\text{Eta}^2$ , \*\* $P$ -values $< 0.001$ , \* $P$ -values $< 0.05$ (GLM).

The interaction of three variables, maternal age groups, sex of newborn and parity was analyzed in another statistical model. Patients were divided into three age categories ≤19 years (teens age), 20–39 years and ≥ 40 years (advanced maternal age) by maternal age at conception in this analysis. These age categories were also used in similar studies focused on the teenaged women [3, 11] and on the advanced maternal age [10, 26]. No covariates were included in this basic model. Table 3 shows results (*P*-values) of statistical analyses (GLM) and Fig. 2 shows direction and size of the effects.

**Table 3.** Effects of age, parity, sex, and their interaction on pregnancy output variables.

|  | *birth weight* | *birth length* | *weight gain* | *pregnancy length* |
| --- | --- | --- | --- | --- |
| age | < 0.001 | < 0.001 | < 0.001 | < 0.001 |
| parity | 0.428 | 0.323 | 0.251 | 0.142 |
| sex | 0.037 | 0.047 | 0.360 | 0.435 |
| age*parity | 0.894 | 0.838 | 0.153 | 0.512 |
| age*sex | 0.195 | 0.056 | 0.904 | 0.101 |
| parity*sex | 0.289 | 0.061 | 0.271 | 0.006 |
| age*parity*sex | 0.171 | 0.068 | 0.221 | 0.019 |

Maternal age, parity, fetal sex and adverse fetal outcomes

**Fig 2.**
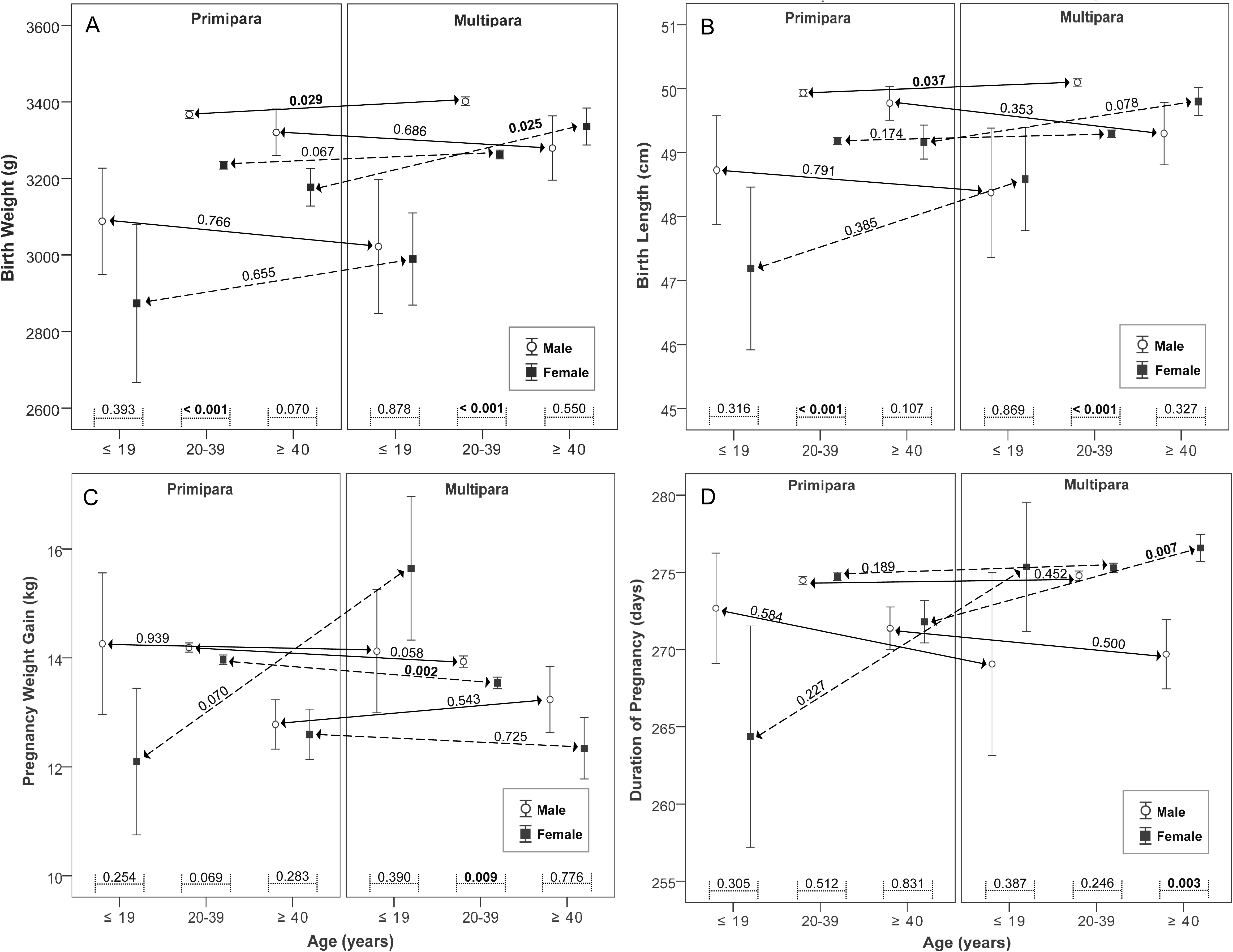
Comparison of the birth weight (A), birth length (B), pregnancy weight gain (C) and duration of pregnancy (D) in interaction women with three different age groups, fetal sex and parity. Left graphs show primiparous and right graphs multiparous women. Full squares show women with female newborn and unfilled circles show women with male newborn. The x-axis shows the age groups by maternal age at conception: 1: ≤ 19 years, 2: 20–39 years and 3: ≥40 years. Vertical bars denote +/- standard errors. The numbers above the lines between the bars show *P*-value of statistical analyses (t-test).

Because of non-monotonic effect of maternal age, this variable was entered into the model as the categorical variable (three age categories ≤ 19 years, 20–39 years and ≥ 40 years), sex (sex of newborn) and parity (primiparous /multiparous women), the values in the table show *P-* values of statistical analyses (GLM).

In our data, we observed 7.9 % of preterm deliveries (before 37 weeks), 6.2 % low birth weight rate (< 2500g), 1 % macrosomia > 4,500g and 32.7 % caesarean deliveries in our data set. Table 4 shows the influence of the maternal age on the binary variables (0/1): preterm delivery, low birth weight, macrosomia > 4,500g and caesarean delivery, separately for primiparous resp. multiparous women and women with male resp. female newborns. The age limits for the comparison of risks of adverse delivery outcomes were determined based on the results of previous studies focused on the study of similar variables [7, 11, 27, 28]. The odds of preterm delivery, low birth weight, macrosomia > 4,500g and caesarean delivery were statistically compared in selected age groups for primiparous resp. multiparous women and women with male resp. female sex of newborns (Table 5). Women ≤ 19 years had significantly higher rates of preterm delivery than women 20–39 years (*P* = 0.012, OR = 3.62, CI_95_ = 1.32-9.90) in primiparous with female newborns. Women older than 40 years had significantly higher rates of preterm delivery than women 20–39 years old (*P* = 0.003, OR = 1.86, CI_95_ = 1.23-2.82) in women with male newborns. To assess the influence of maternal age on the low birth weight, the women were subdivided into two age groups: maternal age 24 years and maternal age ≥ 24 years. The younger women expressed higher rates of low birth weight (*P* < 0.001, OR = 1.87, CI_95_ = 1.44-2.42). This effect of age was significant for both the primaparous and the multiparous women, regardless to the sex of newborns. Similarly, the interaction of parity and sex of the newborn was significant in all models. The older (age ≥ 35 years) primiparous women with male newborns expressed higher incidence of macrosomia (*P* = 0.021, OR = 2.04, CI_95_ = 1.12-3.71) compared to other groups of women. The probability of caesarean delivery increased with the age in the 6 age categories (*P* < 0.001, OR = 1.99, CI_95_ = 1.61-2.46) for primaparous and multiparous women, for male and female newborns and in all models. The probability was also significantly affected by the parity-sex of newborn interaction (Table 5).

**Table 4.**
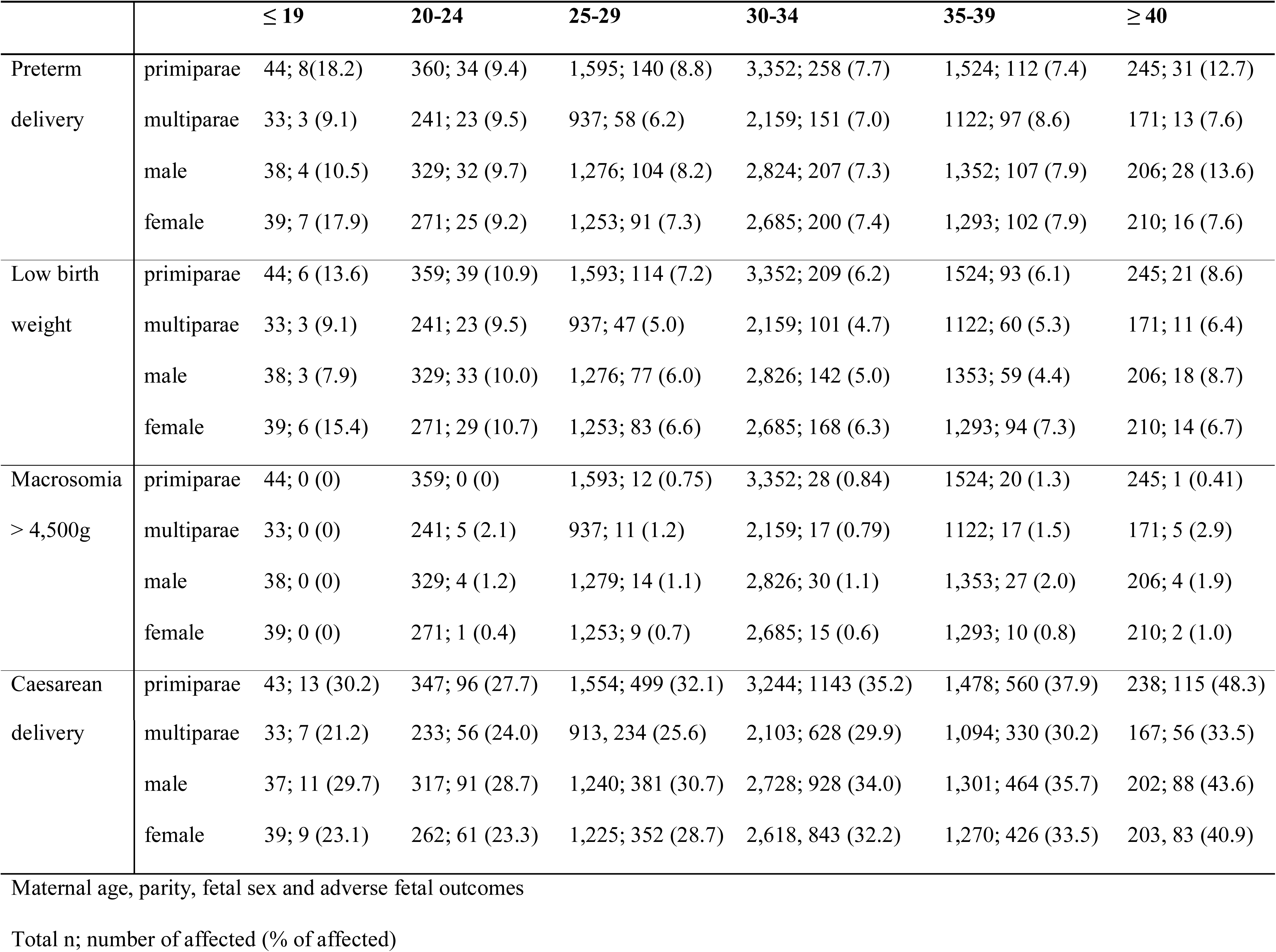
Delivery outcomes in both primiparous/multiparous women and women with male/female sex of the newborn across different age category.

**Table 5.**
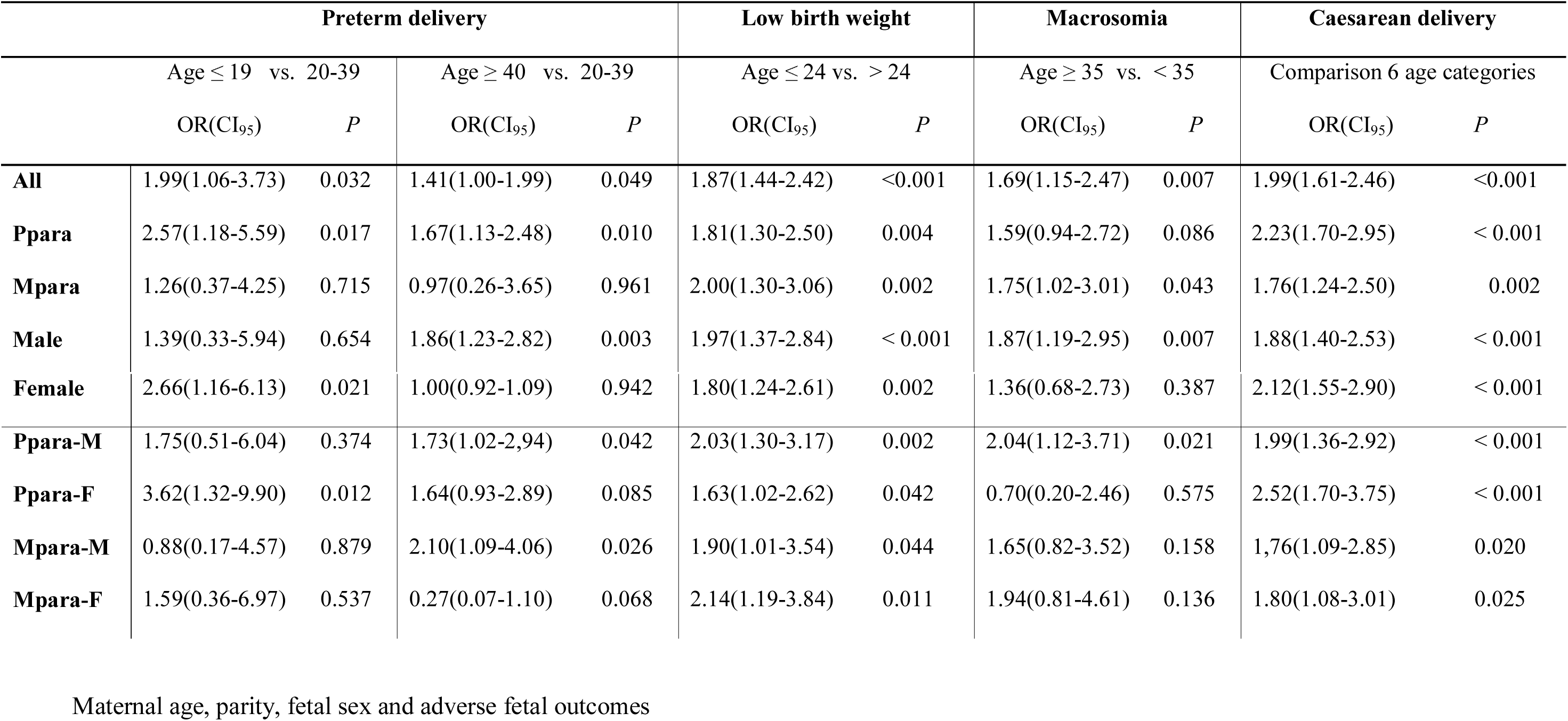
Delivery outcomes in primiparous/multiparous women (Ppara/Mpara), in women with male/female sex of newborn (M/F) and in their interactions, comparisons for different maternal age category, logistic regression (*P*-values), Odds ratio (OR), 95% confidence interval (CI_95_)Maternal age, parity, fetal sex and adverse fetal outcomes.

## Discussion

The results of our study have confirmed that maternal age at conception significantly affects the perinatal outcomes. The observed functions of age were always non-linear. With increasing age, the birth weight and birth length of the newborn increased up to maternal age of 35–39 years, and after this age these rates decreased again. We saw a similar trend for the variable duration of pregnancy, however, duration of pregnancy increased up to maternal age of 25–34 and after this age these rates decreased again. The maternal weight gain during pregnancy decreased from the maternal age of 20–24 years. This non-linear age effect on the variables is in agreement with already published data. Women who are too young [5, 11–14] or older women [4, 6, 10] are a high risk group of women with more frequent perinatal complications. It is known that weight gain during pregnancy and duration of pregnancy are affected by the maternal weight before pregnancy [29, 30] and also by interaction of the maternal Rh-factor and latent infection with protozoan *Toxopalsma gondii* [31]. In our study, the effect of maternal age on weight gain during pregnancy and on the duration of pregnancy was relatively small, however, it was comparable with the effect of maternal weight before pregnancy on these variables.

The birth outcomes were affected by the parity and sex of the newborns in all age groups. Mothers aged 20–39 gave birth to significantly heavier and longer boys than girls and parity resulted in no effect on sex of the newborns. These results are in accordance with previous studies, which found that boys weighed at birth more than girls [17, 18]. In the age group of 20–39 years, the primiparous women also gained more than the multiparous women, with no difference in the sex of the offspring (near the formal border of significance). In addition, mothers with male fetuses gained more weight than mothers with female fetuses, both in primiparous (near the formal border of significance) and multiparous, which is in accordance with the higher birth rates in boys in this age group. In the age categories of younger (≤ 19) and older (≥ 40) mothers, these effects were not likely to be observed for reasons of low N and greater variance of data in these age categories. However, visual inspection of data (Fig. 2) suggests that maternal age alone has a strong influence on birth weight, while the sex of newborns doesn’t play a role in these age groups. It must be emphasized that the previous studies showing higher birth rates in boys were performed in mothers without taking into account their age.

In mothers aged 40 and more, we found higher birth weight and length in newborns of multiparous than in primiparous women, both for girls (birth length was near the formal border of significance) and for boys (significant). This may be related to the fact that the incidence of macrosomia was observed more frequently in male newborns in the older multiparous mothers (≥35) in our data set. Higher delivery rates for girls could then be due to a longer pregnancy, as the length of pregnancy was also affected by the parity and sex of the newborns in mothers aged 40 years and more. The multiparous who gave birth to a girl had the longer pregnancy in comparison to primiparous who gave birth to a girl. Longer pregnancies were also recorded in multiparous mothers with girls than in multiparous mothers who gave birth to a boy.

Significantly more preterm births were observed in very young women and among women of advanced maternal age. The preterm birth was recorded twice as often in young mothers (≤ 19) than in mothers aged 20–39. In addition, 2.5 times more often in the group of primiparous and up to 3.5 times more often in the group of primiparous who gave birth to a girl. Several other studies have reported more premature newborns among parents of early age [11, 28, 32]. The incidence of preterm delivery can be affected by social factors such as the low socio-economic status of the family, the low level of medical care during pregnancy, maternal stress, and the hostile environment at the place of residence [33–35]. These women tend to be in poor health, and according to the T-W Hypothesis [36], they could be more likely to give birth to girls than boys. This could explain our observation of the effect of maternal age on the preterm birth in young primiparous mothers of girls.

The preterm birth was also recorded 1.5 times more frequently in the older primiparous (≥ 40) than in primiparous in the 20–39 aged mothers and almost twice as often in older mothers (≥ 40) with a boy than in mothers aged 20–39 with a boy. The preterm delivery was 2 times more common in multiparous women who gave birth to a boy and the likelihood of preterm birth decreased in multiparous women who gave birth to a girl in comparison to the control group (age 20–39). These findings are largely consistent with those of other relevant studies showing that women of advanced maternal age are at greater risk for having preterm birth [10, 27]. A higher incidence of preterm birth has been observed among mothers of male newborns compared with mothers of females [17], however, no work has documented the combination of dependence between sex, parity, and age.

Maternal age at delivery was significantly higher for macrosomic neonates [7], maternal age ranging between 30 and 39 years, multiparity and gestational age ≥40 years were significantly associated with fetal macrosomia [4]. In our data set we have confirmed a more frequent occurrence of macrosomic neonates in multiparous in older age (≥ 35). An increased risk for macrosomic neonates was also observed in older mothers (≥ 35) who gave birth to a boy, and this effect was strongest in primiparous (OR = 2). Also, previous studies demonstrated that women carrying male fetuses are at increased risk of macrosomia [7, 37, 38]. Although macrosomia is associated with a higher gestational age, it has been shown in our data set that older mothers (≥ 40) carrying male fetuses have a higher risk of preterm birth, which is inconsistent with current studies [4]. Based on our analysis, primiparous women aged ≥ 35 carrying the male fetus are the riskiest group of mothers associated with fetal macrosomia.

In our study, the low birth weight of the newborns was observed almost twice as often in all young women (24 ≤) than in women over the age of 24, regardless of the parity and sex of the newborns. Our results are in accordance with studies that have shown that teenage groups (<25) were associated with increased risks for low birth weight [11]. Also, pregnant women aged <27 years or >32 years carried a greater risk for having a low birth weight infant compared to those aged 27 years [5]. Our data also showed that the low birth weight was not affected by the sex of the newborns or by the maternal parity as opposed to other adverse birth outcomes.

The risk of delivery by caesarean section significantly increased with the maternal age, regardless of the parity and sex of the newborns. Almost 50 % of primiparous mothers older than 40 had caesarean delivery. Based on Table 4, the risk of caesarean delivery was higher in primiparous than multiparous, and also in mothers carrying male fetuses than in mothers carrying female fetuses in all age groups. Our results correspond with previous studies that show an increased risk of caesarean section in women ≥40 [8, 9] and in women carrying male fetuses [18]. Also, the risk of operative deliveries increased with maternal age ≥40 and in pregnancies with male fetuses [26]. A portion of the results could be explained by a more frequent occurrence of neonatal macrosome in mothers aged ≥ 35, as macrosomia is associated with more frequent births by caesarean section [39, 40].

An important limitation of the study was a relatively smaller sample size in sub-analyses evaluated across the different age categories for sex of newborns and both primiparous and multiparous women. The second limitation of the study was the absence of information about maternal socio-economic status (such as prenatal care, marital status, residence, educational level, tobacco and alcohol consumption). The great benefit of this study is that it has been done across all age categories and was not limited to the selected age group of women as in most studies, and different risk age limits could be set based on observed values.

## Conclusions

The results of our study could facilitate risk assessment and consequent optimal health care for pregnant mothers in high risk age categories with respect to parity and possible knowledge of the sex of the child during pregnancy. All results of our research together indicate that “optimal” maternal age without perinatal complications is 25–34 years. The older age is associated with increased risk of complications especially in primiparous women bearing a male fetus. Whereas the pregnancies of younger women bearing a female fetus are at higher risk, too. Our results open further possibilities for research that should analyze the wider spectrum of possible causes of adverse fetal outcomes in more detail.

## Acknowledgments

The study was supported by Charles University Research Centre program No. 204056 and RVO-VFN64165 from the Ministry of Health of the Czech Republic.

